# Characteristics of circadian rhythm-related genes and establishment of a prognostic scoring system (CRscore) for lung adenocarcinoma with experimental verification

**DOI:** 10.1101/2022.07.04.498733

**Authors:** Fengbin Zhang, Qi Zhang, Taie Jiang, Xiaoyu Fan, Wenmiao Wang, Lehan Liu, Kun Liu

## Abstract

**Background:** Non-small-cell lung cancer (NSCLC) is one of the most common malignant tumors worldwide. Lung adenocarcinoma (LUAD), which is the m ain subtype of NSCLC, has a poor prognosis. In recent years, circadian rhythm (CR)-related genes (CRRGs) have demonstrated associations with tumor occurrence and development, but the relationship between CRRGs and LUAD is not clear.

**Methods:** Based on data from The Cancer Genome Atlas and Gene Expression Omnibus databases, we explored the biological function and immune cell infiltration of LUAD in different CR clusters and quantified CR genes using principal component analysis. Then, we built a CR scoring system (CRscore) to explore the relationship between CRRGs and LUAD prognosis.

**Results:** Patients were divided into three clusters (A, B, and C). Biological characteristics, such as survival, immune cell infiltration, and gene enrichment, were significantly different among the three clusters. We then established the usefulness of the CR score, which could effectively predict the prognosis of LUAD. Specifically, patients with a high CR score had a better prognosis and were more sensitive to chemotherapy than patients with a low CR score.

**Conclusion:** CRRGs can be used to assess the prognosis of patients with LUAD. Quantification of CR using the CRscore tool in patients with LUAD could help to guide personalized cancer immunotherapy strategies in the future. Thus, the CRscore may be a powerful prognostic tool for LUAD.

## Introduction

In recent years, circadian rhythm (CR)-related genes (CRRGs) have become a hot topic in cancer research. Many studies have shown that CRRGs regulate cell proliferation, malignant tumor cell apoptosis, and neuroendocrine and immune function. CRRGs are expressed in many behaviors and physiological processes, including tumor occurrence and development (1) . Disruption to the CR plays a key role in tumorigenesis and promotes the establishment of cancer features. Moreover, tumorigenesis impairs the CR directly (2). In recent years, increasing attention has been paid to studying the effects of the CR in the human body. For example, its role in tumorigenesis, cancer characteristics, treatment options, and how CRRGs work are becoming interesting topics for future research (3, 4). There are strong links between cancer and CR disorders. For example, transcription of core CRRGs affects the efficacy of treatment and the prognosis of a variety of cancers (5–7). However, the mechanisms regulating the effects of CRRGs on clinical prognosis remain unclear.

A previous study showed that disruption of the CR can promote the development of lung tumors (8). Lung cancer is the leading cause of cancer death, accounting for 18.4% of all cancer deaths, and has the highest incidence of all types of cancer worldwide (11.6%) (9). Non-small-cell lung cancer (NSCLC) is the most common type of lung cancer, accounting for approximately 85% of cases. Lung adenocarcinoma (LUAD) is the most common type of NSCLC, and its incidence is increasing year by year. Therefore, it is necessary to identify key molecules and to establish an effective prediction model with good stability that can be used to implement precise treatment and improve the prognosis of patients with LUAD. In this study, we investigated the relationship between LUAD prognosis and CRRGs and established a CR scoring system (CRscore) to predict the prognosis of patients with LUAD and guide treatment selection.

## Materials and methods

### Data source and pre-processing

The data and clinical information of patients with LUAD and somatic mutation data were obtained from The Cancer Genome Atlas (TCGA). In addition, another set of files (GSE37745) was downloaded from the Gene Expression Omnibus (GEO) database to ensure the adequacy of the sample size. Subsequently, in UCSC Xena (http://xena.ucsc), we downloaded the copy number data of LUAD (10). Using the limma package, the standard human gene expression matrix of each independent sample was converted from the TCGA gene expression profile, and Transcripts Per Million data were converted from Fragments Per Kilobase Million data (11). The standard human gene names were converted from two GEO files in Perl, as was the clinical information obtained from the GEO. The batch effects of the data were corrected using the sva package of R (12). We combined the statistical data of TCGA-LUAD and GSE37745 as the combined queue. We used R (version 4.1.1) and Strawberry Perl (version 5.32.1) to process the data.

### Differential expression of CRRGs

This study examined 10 genes (AANAT, NPAS2, ARNTL, CRY1, PER3, CLOCK, CRY2, CSNK1E, NR1D2, and BHLHE40). First, the copy numbers of the CRRGs were extracted from TCGA-LUAD using Perl software, and histograms were intuitively constructed using R software. The R Circos package was used to map the change in 10 CRRGs on 23 pairs of chromosomes to investigate the relationships between CRRG copy number and chromosome. The differential expression of these CRRGs in TCGA-LUAD was compared using the Wilcoxon rank-sum test in the limma package, which provides a comprehensive solution for microarray and RNA-Seq differential analyses (13). A waterfall plot was created using the maftools package of R to bear out the mutation rate of CRRGs in patients with LUAD. A boxplot was constructed using the ggpubr package of R, and a heatmap was drawn using the pheatmap package. A P value of <0.05 was considered statistically significant.

### CR regulator analysis

#### CR clustering

To bear out the value of CRRGs, we used an unsupervised cluster analysis to organize the amalgamated dataset according to the expression of CRRGs. The samples were clustered using the ConsensusClusterPlus package according to the expression of CRRGs. All samples were divided into K=[2-9] groups, and the most suitable CR regulator cluster was obtained according to three conditions after the cycle, including a close connection within types and an unclose connection between types, the number of samples in each cluster was not short, and no significant increase in the cumulative distribution curve area . According to the correlations between the CRRG clusters and the survival status, the survminer package was used to determine the cut-off points of each subgroup of data, and all possible cut-off points were tested to identify the maximum rank statistics. Based on the log-rank statistics, patients were divided into high, medium, and low expression groups. The Kaplan–Meier method and the survminer package were used to generate survival curves for the predictive analysis. The log-rank test was used to identify significant differences. A P value of <0.05 was considered statistically significant.

#### Single-sample GSEA analysis and GSVA analysis

GSVA is a non-parametric, unsupervised method for estimating path changes and changes in the activity of biological processes in samples in an experimental dataset (14). Based on differences in CRRGs, we revealed the biological pathways between different CRRG clusters. For the gene enrichment analysis, we download “C2.CP.KEGG.7.5.1.symbols ”. The scores of different paths in each sample were calculated using the GSA package of R, and the path differences were analyzed using the limma package. A P value of <0.05 indicated differential expression of pathways in pathway regulation (15, 16). Heatmaps were drawn using the phatmap package. We used the single-sample GSEA and the gene enrichment score to inspect the relative abundance of immune cell infiltration to obtain the immune score (17). We obtained the gene sets of every type of tumor microenvironment (TME)-infiltrating immune cell from a previous study (18), including activated dendritic cells, CD8+ T cells, regulatory T cells, and macrophages. The pertinence of CRRG clusters and immune scores was explored using the limma package, and the ggpubr package was used to draw the box graph. A P value of <0.05 was considered statistically significant.

#### Differential analysis

To identify differentially expressed genes (DEGs) related to CR, we used the limma and VennDiagram packages to identify DEGs between CRRG clusters. DEGs with an adjusted P value of <0.001 were reserved (19). Gene Ontology (GO) and Kyoto Encyclopedia of Genes and Genomes (KEGG) analyses were used to examine the pathways and functions of DEGs. P values and Q values of <0.05 were used to identify the potential pathways and biological functions of these DEGs . According to the number of enriched CRRGs, we chose the top 30 KEGG pathways and GO pathways.

### CRRG clusters

To identify the CRRGs there were associated with prognosis (P < 0.05), the univariate Cox regression analysis was used to analyze the DEGs using the survminer package. The ConsensusClusterPlus package was used to cluster the samples according to the expression of prognostic CRRGs to determine the CRRG clusters that should be further analyzed. First, we performed a survival analysis to evaluate the prognostic value of the CRRG clusters using the survminer package. Then, the patients were divided into three groups: A, B, and C. The Kaplan–Meier method was used to draw the survival curves of the three groups, and the difference between the three groups was significant (P < 0.001) according to the log-rank test . After collecting the clinical data (LUAD stage, age, sex, and alive/deceased status), the phatmap package of R was used to draw a heatmap of the correlations between clinical features and CRRG clusters. The boxplot was drawn using the ggpubr package.

### CR score

To quantify the expression of CRRGs in patients with LUAD, we constructed a CR scoring system (CRscore) based on the CRRGs associated with prognosis. Then, we used the principal component analysis (PCA) to construct the CRscore. The PCA can effectually identify the most significant portions and structures in the data, eliminate redundancy and noise, reduce the dimension of primordial intricate data, and uncover the simple structure hidden behind the mazy data.

We used the following calculation to construct the CRscore mainly using the PCA:

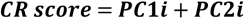

where i represents the expression of prognostic DEGs.

### Correlations between the CR score and patients’ clinical characteristics

We divided the patients into the high CR score group and the low CR score group for further analysis using the CRscore. First, the survival analysis was used to assess the prognostic value of the CRRG clusters using the same method as described above. Then, we analyzed the relationships between the CRscore and patients’ clinical features, including age, sex survival status, and LUAD stage, to identify the relationship between the CR score and survival in the context of the different individual clinical characteristics using the univariate and multivariate Cox regression analyses. Patients with the same clinical characteristics were analyzed respectively to exclude the influence of clinical characteristics on the conclusion . Then, the correlations between CR clusters, gene clusters, CR grouping, and clinical data (LUAD stage, age, sex, alive/deceased status) were measured using the ggalluvial package for Mulberry plots. In addition, differences in the CR score were calculated for the different clinical characteristics (stage, age, and sex). The plyr and ggpubr packages were used to construct percentage plots and box-line plots, respectively. The CR score and CR stage were compared using the limma package. The log-rank test was used to identify statistically significant differences, and the Kaplan–Meier method was used to analyze each clinical feature based on the CR score.

### Correlations between the CR score and the tumor mutation burden (TMB)

The TMB is the sum of somatic gene-coding errors, gene insertions, deletion errors, and base substitutions per million bases. First, using Perl software, the TMB was calculated for each sample. A correlation diagram and boxplot of the relationship between the TMB and CR grouping was constructed using the ggpubr package of R. Then, the survminer package was used to perform the survival analysis. According to the TMB, all samples were divided into two groups: the low expression group and the high expression group. We also used the Kaplan–Meier method to plot the survival curves based on the TMB and the TMB combined with the CR score. A P value of <0.05 was considered statistically significant with the log-rank test.

### Analysis of immune checkpoint genes

First, the corrplot package was applied to contrast the correlation between the immune score and the CR score. Then, the samples (by CR group) were crossed with the clinical information samples (survival status), and the data were combined using R software. Immunotherapy score files were acquired from the Cancer Immunome Atlas (TCIA) website (https://tcia.at/home). A violin plot was created to observe the relationship between immune checkpoint genes and groups with high and low CR scores using the ggpubr package. We analyzed the relationships between the CR score and the expression of common immune checkpoint (PD-L1, PD-L2, PD1, CTLA4) using the limma package.

### Prognostic treatment of LUAD based on the CR score

The immunophenotypic scores (IPSs) of TCGA-LUAD patients were obtained from the TCIA database (20). The difference in the IPS between the high CR score group and the low CR score group was analyzed to comprehend the immunogenicity of the two groups of patients. We used the pRRophetic package of R to predict the half-maximum inhibitory concentration of five chemotherapeutic agents for the treatment of LUAD, including cisplatin, gemcitabine, paclitaxel, vinorelbine, and methotrexate (21). These five kinds of chemotherapy drugs in LUAD patients with the sensitivity of the forecast are based on the cancer drug sensitivity genomics (GDSC, https://www.cancerrxgene.org/).

### Statistical analysis

The Kruskal–Wallis test was used to compare three or more groups. Using the Surv-Cutpoint function in the survminer package of R, the patients were divided into two groups: the high CR score group and the low CR score group. Spearman’s and distance correlation analyses were used to calculate the correlation coefficients between the TME-infiltrated immune cells and CRRG expression. We used Pearson’s correlation to calculate the correlation coefficients between CRRGs. A P value of <0.001 was considered statistically significant. The univariate Cox regression analysis was used to calculate the hazards ratios of the CRRGs and DEGs. The cor.mtest function was used to calculate the relationship between the CR score and immune cell infiltration, and the corrplot package of R was used to visualize this relationship. Spearman’s correlation analysis was used to obtain the correlation coefficient between the TMB and the CR score. The R Circos package was used to show the change in copy number on 23 chromosomes based on the CRRGs identified from TCGA-LUAD (22). The maftools package was used to construct a waterfall plot to show the mutation status of TCGA-LUAD. All data were analyzed using R (version 4.1.1) or Perl (version 5.32.1) software (23). The R packages used in this study and their functions are available from the BioConductivity website.

### Clinical validation of CRRGs

#### Quantitative reverse-transcriptase polymerase chain reaction (qRT-PCR)

We extracted total RNA from freshly isolated tissues utilizing TRIZOL reagent (#15596026; Thermo Fischer Scientific, US) (The experiment was reviewed and approved by the Ethics Committee of Nantong University Affiliated Hospital. Approval Number:2021-K077-01.) . Complementary DNA was synthesized from whole RNA using random primers. The PCR primer sequences of NPAS2 were designed as follows: forward: 5′-CGTGTTGGAAAAGGTCATCGG-3′; reverse: 5′-TCCAGTCTTGCTGAATGTCAC-3′. Reverse transcription was performed at 42℃ for 30 minutes, followed by 85℃ for 5 minutes. The PCR conditions included initial denaturation for 10 minutes at 95°C, followed by 40 cycles of 95°C for 20 seconds, 55°C for 30 seconds, 72°C for 30 seconds, 95°C for 1 minute, and 55°C for 30 seconds. NPAS2 mRNA was quantified by qRT-PCR with SYBR Premix ExTaq (Applied Takara Bio, Baori Medical Biotechnology) and normalized to GAPDH as the reference gene.

#### Western blot

Total protein was extracted from tissue, and the protein concentration was determined using the bicinchoninic acid assay (#23225) (The experiment was reviewed and approved by the Ethics Committee of Nantong University Affiliated Hospital. Approval Number:2021-K077-01.). Then, 20 μg protein was loaded into each well and separated by polyacrylamide gel electrophoresis. The protein was then transferred to a polyvinylidene fluoride membrane using the wet transfer method and blocked with 5% skimmed milk at room temperature for 2 hours. Rabbit anti-human NPAS2 (1:2000, PHR3777) and rabbit anti-human GAPDH were incubated overnight at 4°C. After washing, horseradish peroxidase-conjugated horseradish peroxidase-conjugated goat anti-rabbit secondary antibody (1:10,000, #7076) was incubated at 37°C for 2 hours. The membrane was prepared with enhanced chemiluminescence reagent. The average band strength was measured using Image J software (National Institutes of Health, US). The gray value of the target protein was divided by the gray value of GAPDH to calculate the relative protein expression of the target. All antibodies were purchased from Abmart, china.

## Results

### Epigenetic analysis of CR in LUAD samples

Ten CRRGs were examined in this study. As shown in the waterfall figure, 33 of 561 (5.88%) samples had mutations (Fig. 1C). The mutation rates of PER3, CRY2, NPAS2, CRY1, CLOCK, ARNTL, and CSNK1E were 1%, while the mutation rates of BHLHE40, NR1D2, and AANAT were 0%. The copy number variation (CNV) analysis showed a significant increase in the copy numbers of AANAT, NPAS2, ARNTL, CRY1, PER3, CLOCK, and CRY2, while there was a marked decrease in the copy numbers of CSNK1E, NR1D2, and BHLHE40 (Fig. 1D). The circle diagram shows the chromosomal locations with copy number variations (CNVs) in CRRGs (Fig. 1B). LUAD tissues and healthy tissues can be identified by CNVs in chromosomes. To identify the relationship between regulatory factors and epigenetics, we analyzed the expression of CRRGs (Figs. 1A, E). NR1D2, AANAT, CRY1, NPAS2, CSNK1E, PER3, and CRY2 were significantly differentially expressed between LUAD tissues and healthy tissues (P < 0.05) Changes in the expression of CRRGs may vary with copy number. The CRRGs demonstrated specific epigenetic changes in tumor tissues and adjacent non-cancerous tissues. Therefore, seven CRRGs with obvious differences in expression (NR1D2, AANAT, CRY1, NPAS2, CSNK1E, PER3, and CRY2) were studied further.

**Fig 1.**
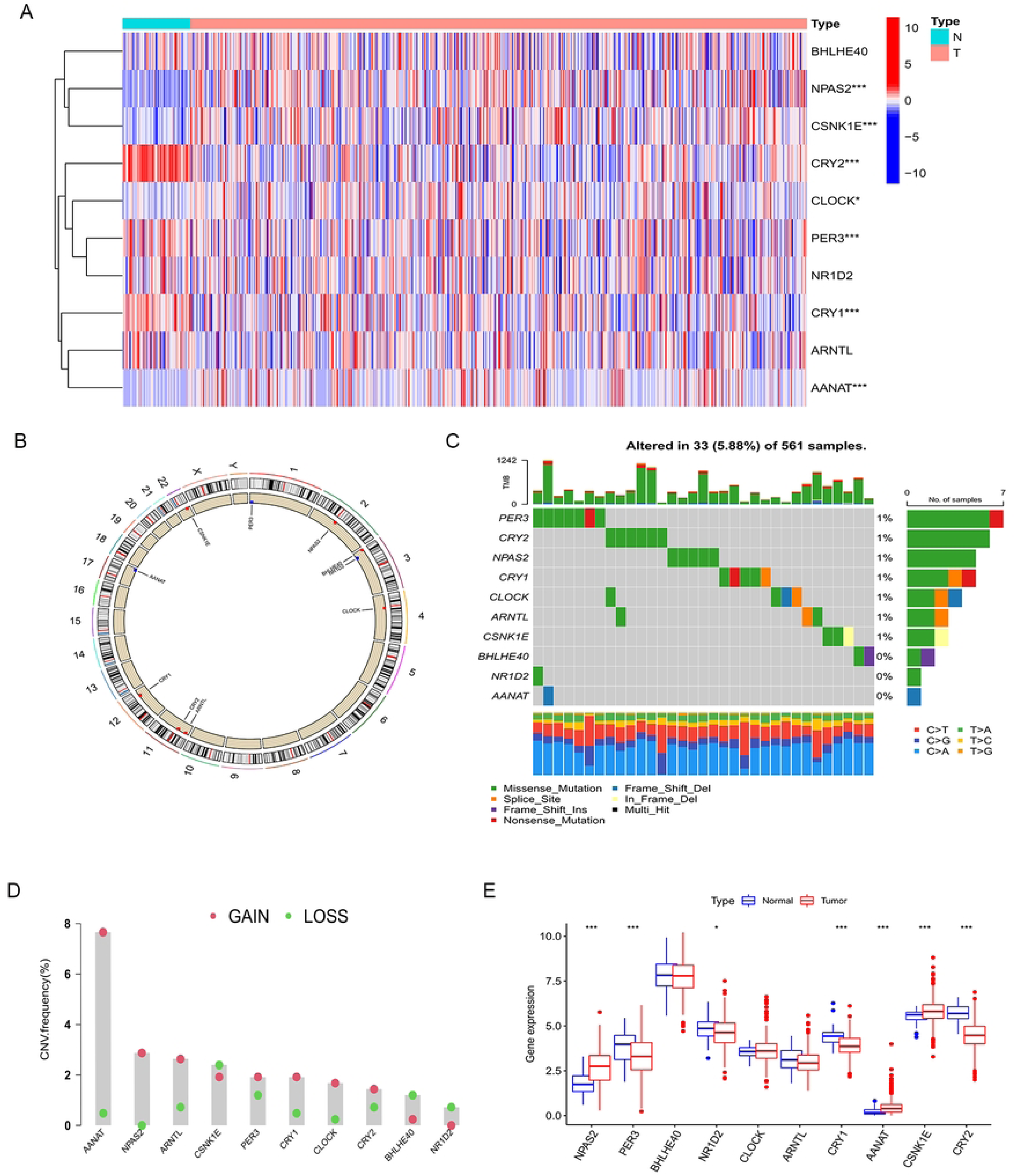
Differences in circadian rhythm genes between LUAD patients and normal patients. **(A)** Heatmap: To clarify the difference expression of circadian rhythm genes between the tumor group and the normal group. **(B)** The change of CNV’s position of circadian rhythm genes was in 23 pairs of chromosomes. **(C)** CRGs waterfall: The number on the right represents the mutation frequency of CRGs in LUAD patients, and the bar chart represents the proportion of mutations per base. **(D)** CNV mutation frequency of CRGs in LUAD: the height of the column represents the mutation frequency. Green dots represent deletions and red dots represent amplifications. **(E)** The difference expression of circadian rhythm genes between normal group and tumor group through boxplot (asterisk indicates statistical P value (* P<0.05; ** P< 0.01; *** P<0.001)

### Unsupervised clustering based on CR

We introduced a new cohort (merged cohort), which consisted of TCGA-LUAD data and GEO data (GSE37745). Unsupervised clustering was used to separate the tumor samples according to the expression of the seven CRRGs mentioned above. According to the cumulative distribution function value, the optimal quantitative cluster was determined as 3 (k = 3). Thus, the tumor samples were divided into three groups (Fig. 2A): CRcluster A, CRcluster B, and CRcluster C. There were significant differences among the three clusters according to the results of the PCA (Fig. 5A). Specifically, CRcluster C had the greatest survival advantage (Fig. 2B). The heatmap shows differences in CRRG expression among the different clusters, and the expression of CRRGs was highest in CRcluster B (Fig. 2C).

**Fig 2.**
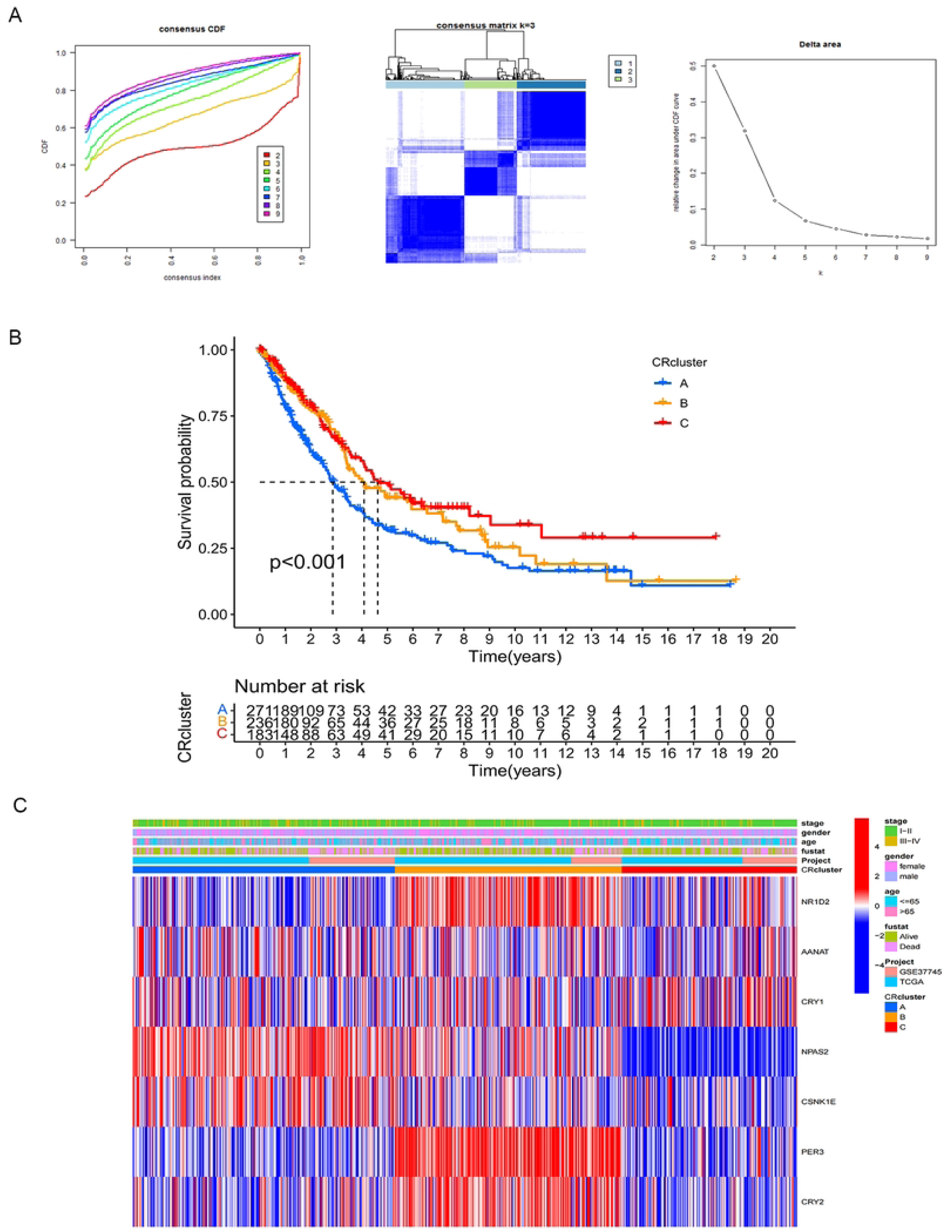
CRGs of unsupervised cluster analysis. **(A)** CRGs uses the method of unsupervised cluster analysis to find that K=3 is the optimal number of clusters. **(B)** Survival analysis of LUAD patients in three different CRclusters. **(C)** seven circadian rhythm genes (NR1D2, AANAT, CRY1, NPAS2, CSNK1E, PER3, CRY2) combined with different clinical characteristics of CRcluster heatmap.

### Differences in immune cell infiltration and function between the CR clusters

To explore the potential biological functions of the three CRRG clusters, the GSVA analysis was performed (Figs. 3A–C). CRcluster A and CRcluster B were mainly related to CR in mammals, such as transforming growth factor-beta, the cellular response to transforming growth factor-beta, and regulation of the transforming growth factor-beta receptor. CRcluster C was related to protein expression, such as the mitogen-activated protein kinase (MAPK) signaling pathway and neurodegeneration. There were significant differences in immune cell infiltration among the three clusters based on the results of the GSEA analysis (Fig. 4C). We found that CRcluster C had a mass of infiltrating immune cells, including CD4+ T cells and CD8+ T cells, which are involved in specific immunity. CRcluster A included dendritic cells, macrophages, and monocytes, which participate in the non-specific immune response. CRcluster B included eosinophils, immature dendritic cells, myeloid-derived suppressor cells, macrophages, mast cells, and plasmacytoid dendritic cells (P < 0.05). Patients in CRcluster C revealed a favorable survival status (Fig. 2B). In combination with the results of the GSVA analysis, we predicted that immune cell infiltration may play a major anti-tumor role (21).

**Fig 3.**
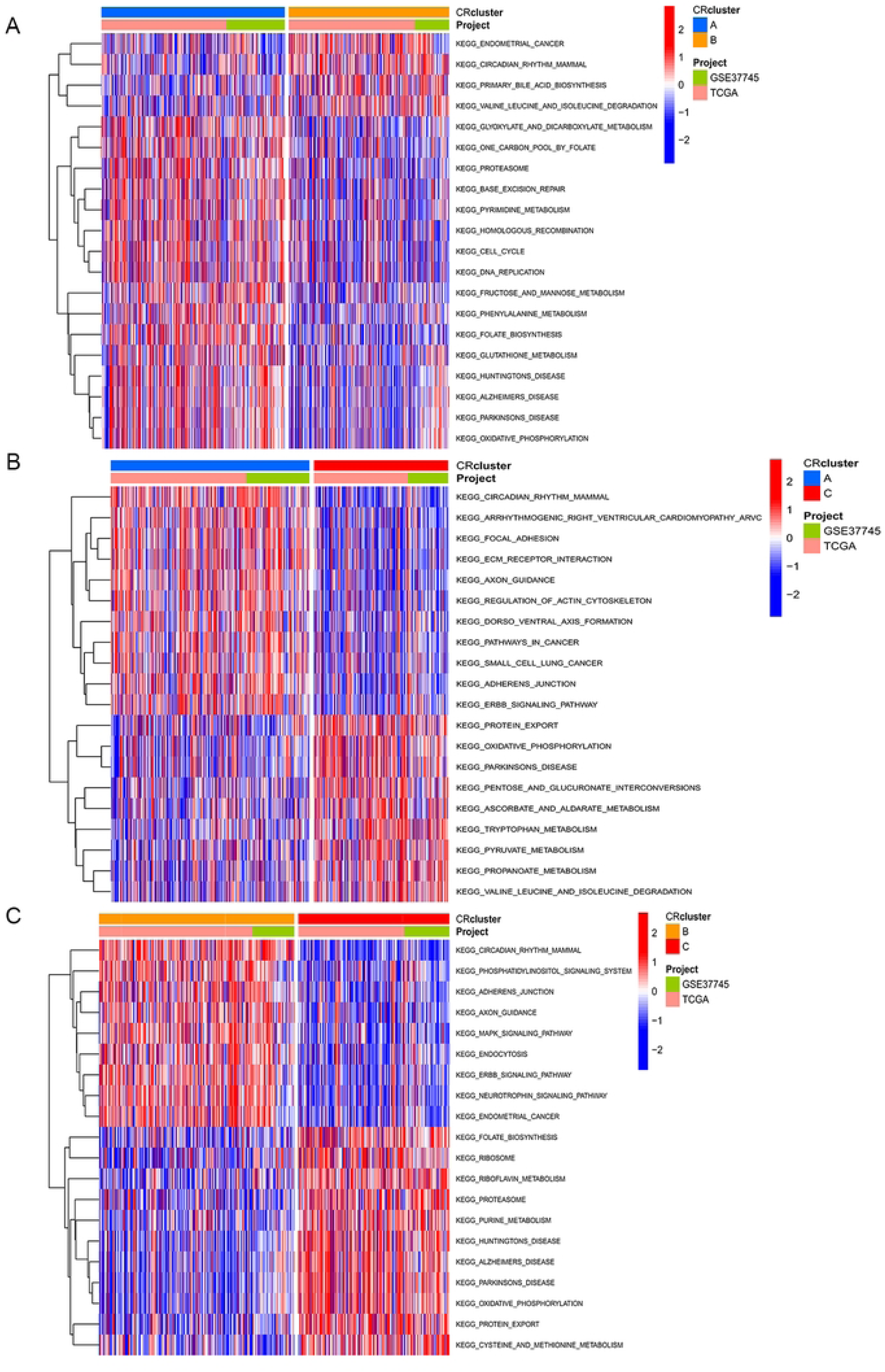
GSVA analysis: the activation status of biological pathway of each CRcluster was observed in pairs in 3 groups of CRclusters. **(A)**clusterA-clusterB **(B)**clusterA-clusterC **(C)**clusterB-clusterC

**Fig 4.**
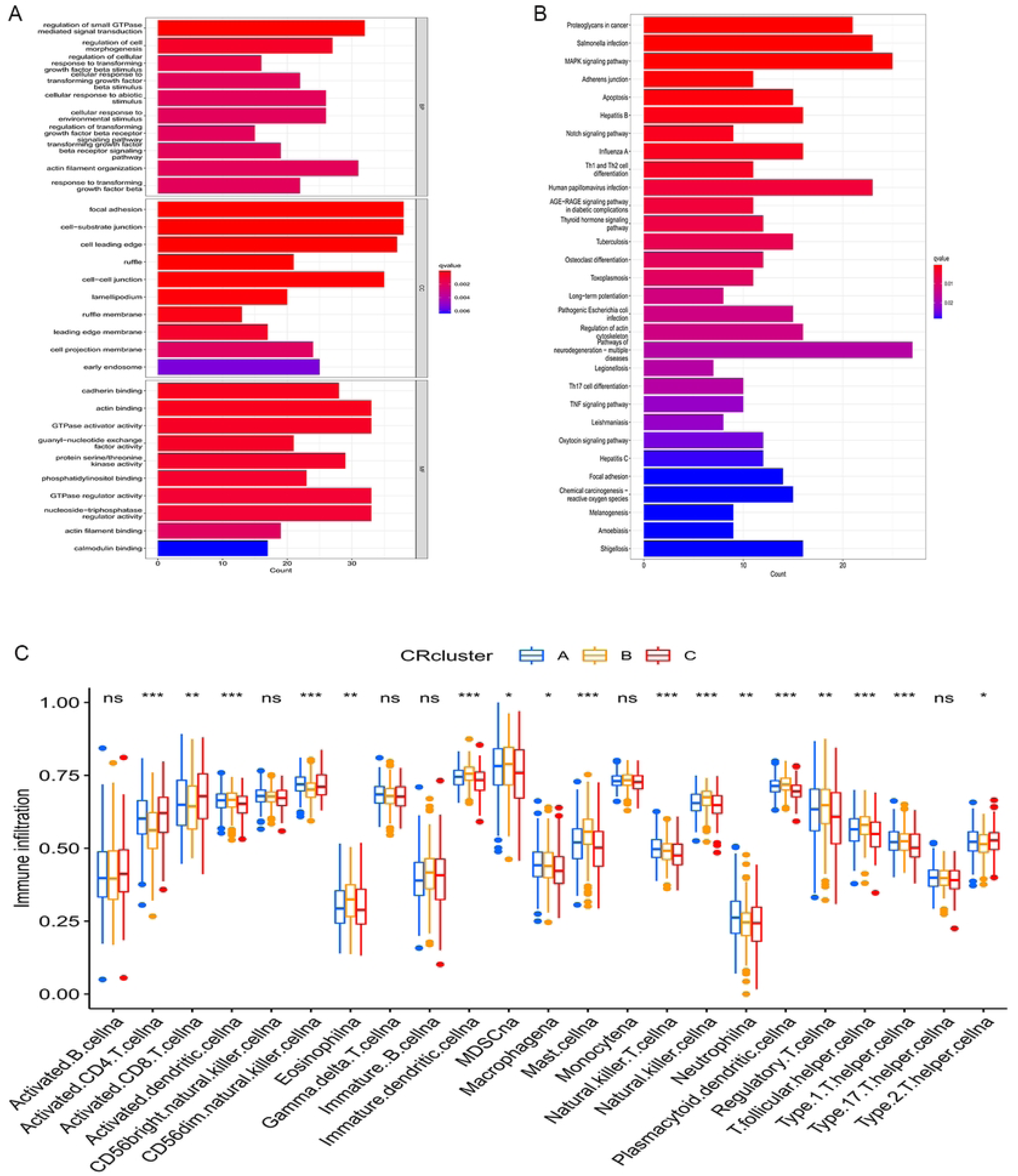
Functional analysis of DEGs. **(A)** GO enrichment analysis of 579 DEGs intersected in three CRclusters. **(B)** KEGG enrichment analysis of 579 DEGs intersected in three CRclusters. **(C)** The abundance of each immune-infiltrating cell in the three CRclusters, the boxline of the boxplot represents the median, the dot outside the box represents the outlier, and the asterisk represents the P value (* P<0.05; ** P< 0.01; *** P<0.001)

**Fig 5.**
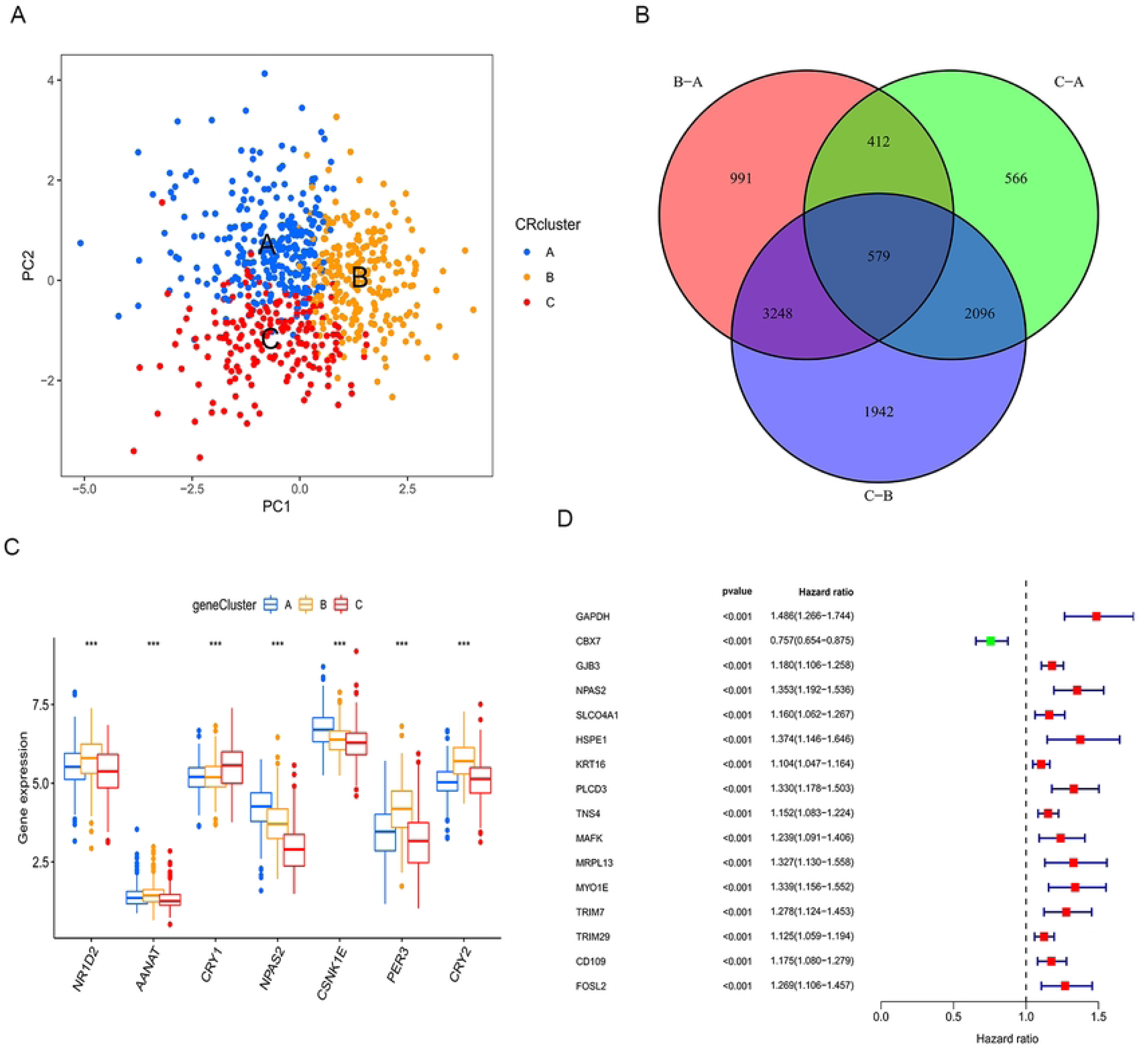
The characteristics of DEGs. **(A)** Principal component analysis(PCA): significant differences in the transcriptome of the three CRclusters. **(B)** The Venn diagram shows 579 differential expressed genes (DEGs) of circadian rhythm-related genes in the three CRclusters. **(C)** Differences in the expression levels of seven circadian rhythm genes in the three CRclusters. **(D)** Forest map: the top 16 genes in the prognostic DEGs.

### Identification of DEGs related to CR

Based on the three CR clusters, we conducted a differential analysis of the amalgamated cohort to identify DEGs related to CR. As shown in Fig. 5B, there were 5,230 DEGs between CRcluster A and CRcluster B; 3,653 DEGs between CRcluster A and CRcluster C; and 7,865 DEGs between CRcluster B and CRcluster C. A total of 579 DEGs related to CR with adjusted P values of <0.001 were selected based on the co-intersection of the CR typing differential genes. These genes were analyzed using GO and KEGG enrichment analyses to identify their functions (Figs. 4A, B). We identified 151 GO terms and 49 KEGG pathways (P < 0.05, Q < 0.05). The top 30 GO terms and KEGG pathways with the greatest number of genes were screened. The results showed that the GO terms and KEGG pathways above were mainly involved in the MAPK signaling pathway, the cellular response to environmental stimuli, the cellular response to abiotic stimuli, protein serine or threonine kinase activity, and melanogenesis. We then performed the univariate Cox regression analysis on the 579 DEGs related to CR and identified 110 DEGs (P < 0.05). We identified 16 prognosis-related DEGs (Fig. 5D), which showed a significant ability to predict patient survival (P < 0.05). Coincidentally, the patients were still divided into three clusters (A–C) using the unsupervised cluster analysis of the 110 CRRGs (Fig. 6A). Patients with gene cluster B had the greatest survival rate in the survival analysis (Fig. 6B). We found that the proportion of patients with LUAD stage I−II was particularly large, and these patients were mainly concentrated in gene cluster B (Fig. 6C). Subsequently, on the basis of the three gene clusters, we analyzed the differential expression of the seven CRRGs in the merged cohort using the limma package. As expected, there were remarkable differences in CRRG expression among the three gene clusters (Fig. 5C). The regulatory mechanism of CRRGs was verified based on the above results.

**Fig 6.**
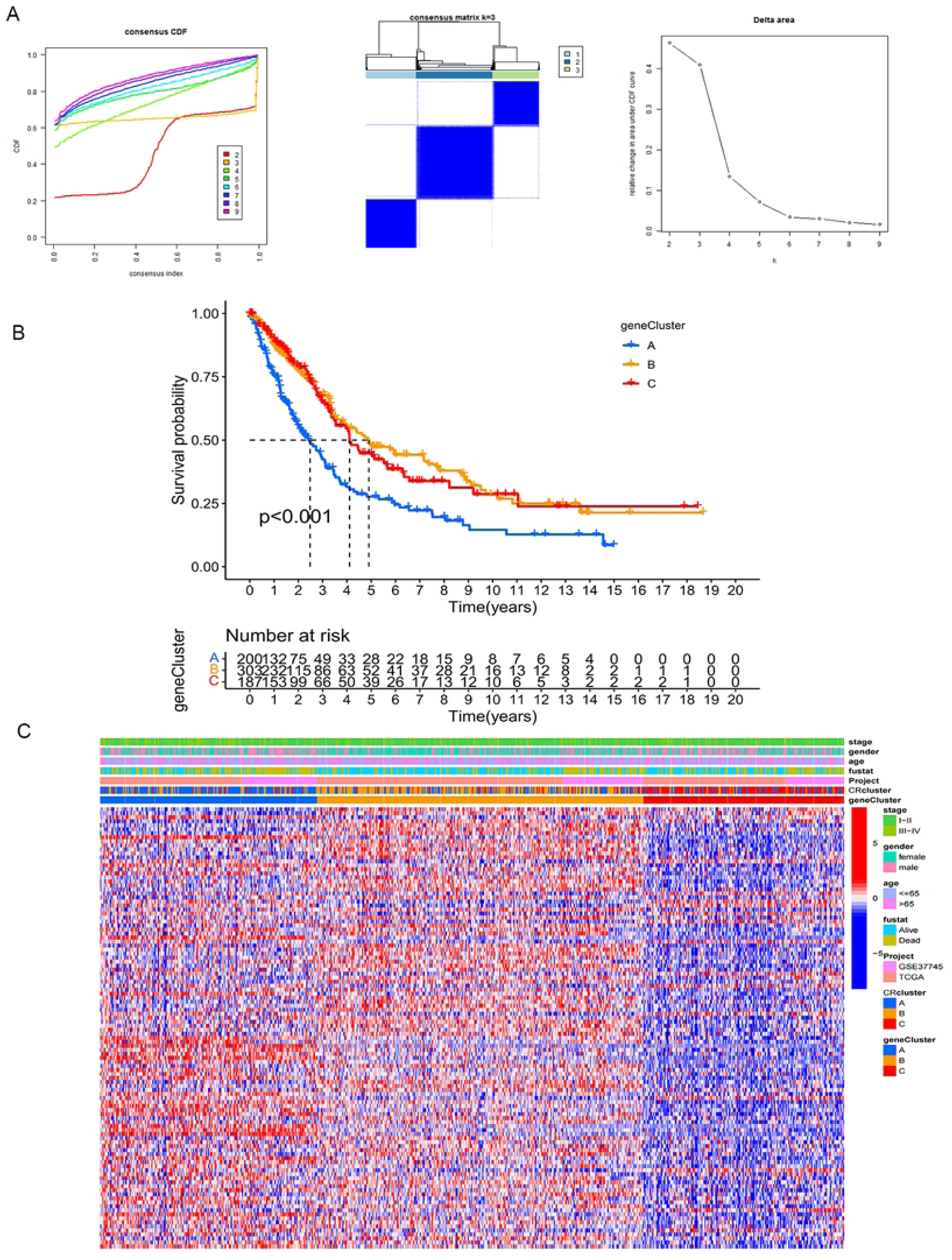
Unsupervised cluster analysis of prognostic associated DEGs. **(A)**Unsupervised cluster analysis was used for prognostic genes in DEGs to determine that K=3 is the optimal cluster number. **(B)** Comparison of the overall survival of the three CRclusters. **(C)** CRcluster heatmap: 110 circadian rhythm-related genes associated with prognosis combined with different clinical characteristics.

### Relationship between the CR score and traits of each subtype

We established a scoring system (CRscore) to quantify the expression of the 110 DEGs related to CR as prognostic predictors. The survival rate of patients with a high CR score was significantly higher than the survival rate of patients with a low CR score according to the survival analysis (Fig. 7A). Changes in the clinical characteristics (LUAD stage, age, sex, alive/deceased status) and subgroups of patients are shown in the Sankey diagram (Fig. 7B). Most immune cells were negatively correlated with the CR score (Fig. 8A), and the infiltrating immune cells were significantly negatively correlated with the CR score (Fig. 8B). In other words, the lower the CR score, the stronger the immunity. The Kruskal–Wallis test showed a significant difference in the CR score between the CR clusters and the gene clusters. CRcluster A had the lowest score, while CRcluster C had the highest score. As a result, we hypothesized that patients with high and low CR scores were more inclined to suppress and develop tumors, respectively. This is well supported by the survival curves of the high and low CR score groups (Fig. 7A). In terms of the gene clusters, the CR score sequence was gene cluster C > gene cluster B > gene cluster A (Fig. 7D). The same was true for the CR clusters (CRcluster C > Crcluster B > Crcluster A) (Fig. 7C).

**Fig 7.**
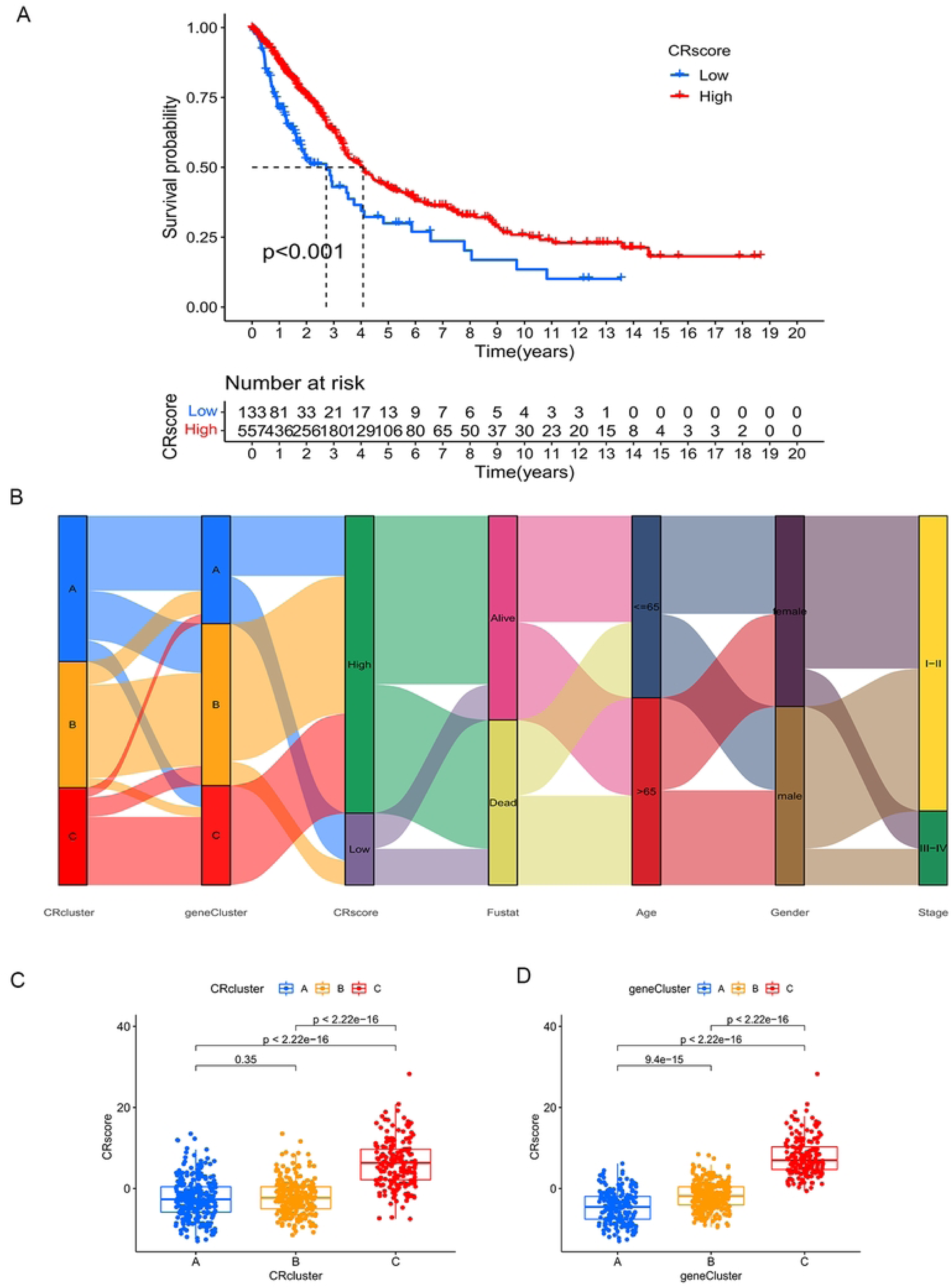
The characteristics of CRscore. **(A)** Comparison of overall survival of high and low CRscore based on circadian rhythm genes. **(B)** The Sankey diagram shows the correlation among CRscore and genecluster, CRcluster, fustat, age, gender, stage. **(C)** Kruskal-wallis test was used to analyze the statistical differences between CRscore and the three CRclusters. **(D)** Statistical differences between CRscore and the three Genecluster (Kruskal-Wallis test analysis, p<0.001)

**Fig 8.**
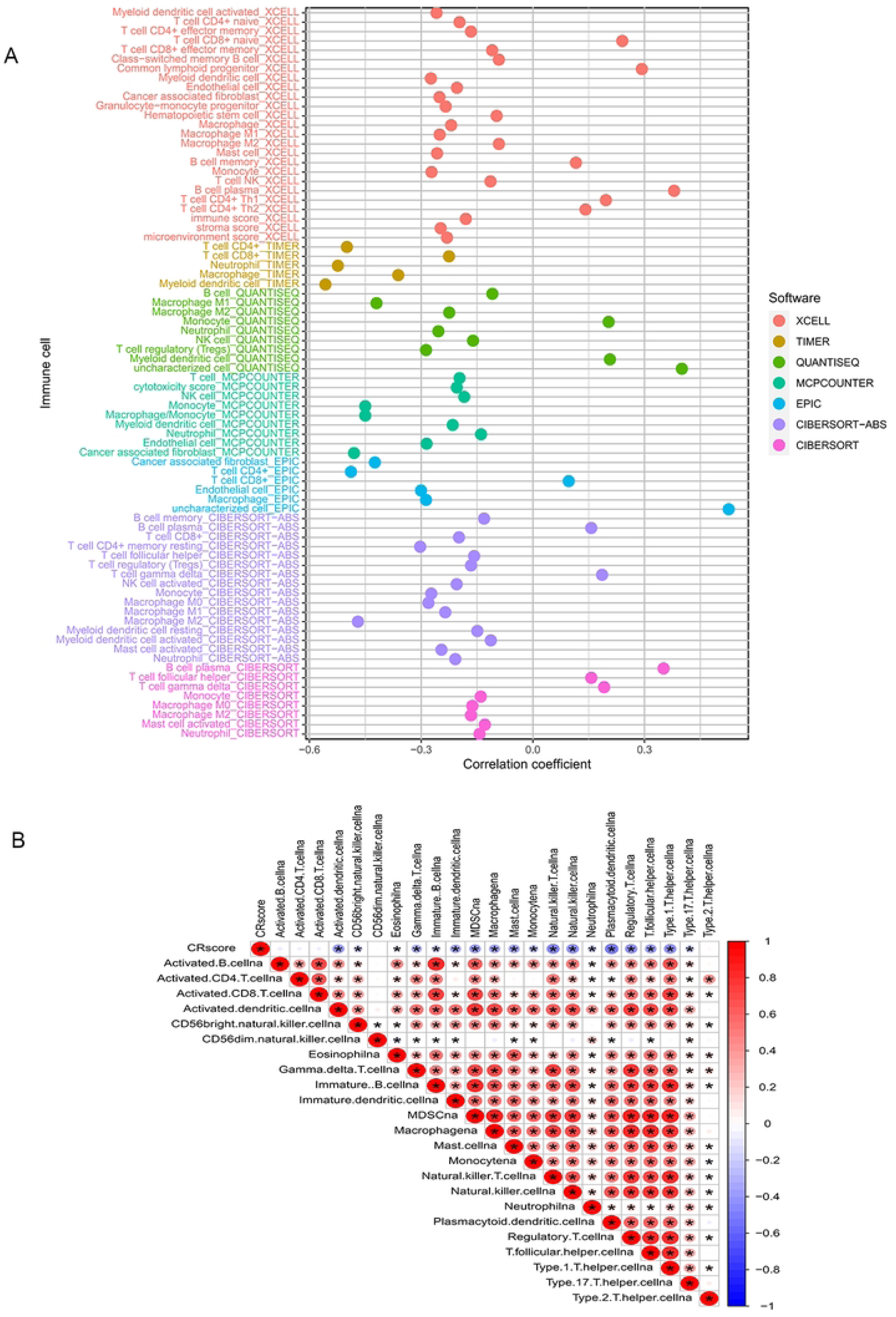
The correlation between CRcsore and immune-infiltrating cells. **(A)** The Correlation between CRcsore and immune-infiltrating cells was detected in seven different software. The Correlation coefficient greater than 0 was positive, and the Correlation coefficient less than 0 was negative. **(B)** Immuno correlation analysis between CRscore and immune-infiltrating cells.

We also analyzed the relationship between the TMB of LUAD and the CR score to explore the relationship between the CR score and tumor occurrence and development. Tumors with a high CR score showed a high TMB (Fig. 9B). In other words, the CR score was positively correlated with the TMB (P < 0.05; r = 0.17) (Fig. 9A). The survival curves of the TMB and CR score show that there was no significant difference in survival between the high and low TMB groups (Fig. 9C), but the TMB combined with the CR score predicted a significant difference in survival (Fig. 9D). Patients with a high TMB and a high CR score had a longer survival time. It can be speculated that combining the CR score with the TMB can enhance the sensitivity of TMB to forecast the effectiveness of immunotherapy in patients with LUAD.

**Fig 9.**
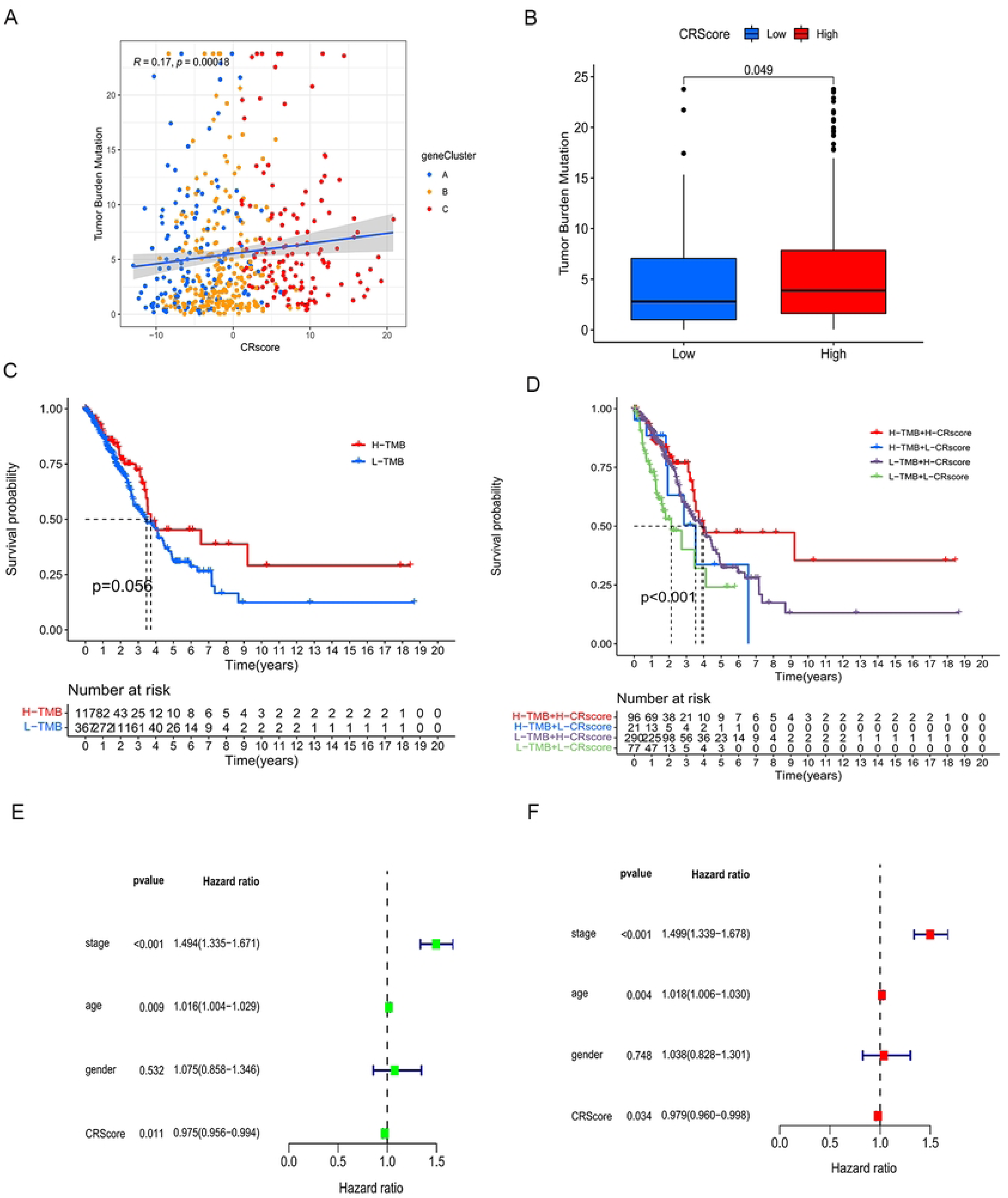
The correlation between CRcsore and TMB. **(A)** The relationship between high and low CRscores and TMB: Scatter plot showed that TMB was positively correlated with CRscore (R=0.17, p<0.001). **(B)** Difference between high and low CRscores and TMB (p<0.05). **(C)** TMB survival analysis: Kaplan-Meier curve was used to describe survival rates of high and low TMB patients. **(D)** CRscore combined with TMB survival analysis: Kaplan-Meier curves were used to depict the survival rates of patients with high and low TMB and high and low CRscore.

### The CR score as a prognostic biomarker

There were no significant differences in the clinical features (with the exception of LUAD stage) and CR score (Figs. 10A–H). Stage I−II LUAD accounted for 82% and 73% of samples in the high and low CR score groups, respectively, while stage III−IV LUAD accounted for 18% and 27% of samples in the high and low CR score groups, respectively (Fig. 10A). The quantitative analysis showed that the CR score was significantly different between stage I−II LUAD and stage III−IV LUAD (P < 0.05) (Fig. 10E). Female patients who were deceased and ≤65 years of age accounted for a large proportion of the high CR score group, while male patients who survived and were >65 years of age accounted for a large proportion of the low CR score group (Figs. 10B–D). According to the quantitative analysis, the CR score of patients with stage I−II LUAD was higher than that of patients with stage III−IV LUAD (Fig. 10A). The survival status of patients in the high and low CR score groups with the same clinical characteristics was analyzed to evaluate the universality of the CRscore tool. By comparing the survival status of the two groups for each clinical feature, we identified that patients with higher CR scores had better survival (Figs. 11A–H). Although the P value of patients with stage III–IV LUAD was >0.05, the usefulness of the CRscore tool in predicting prognosis should not be underestimated (Fig. 11H). Next, the univariate and multivariate Cox regression analyses were performed for the CR score and clinical characteristics (LUAD stage, sex, and age). Both the univariate (Fig. 9E) and multivariate (Fig. 9F) analyses showed that age, stage, and CR score were independent prognostic factors in this study.

**Fig 10.**
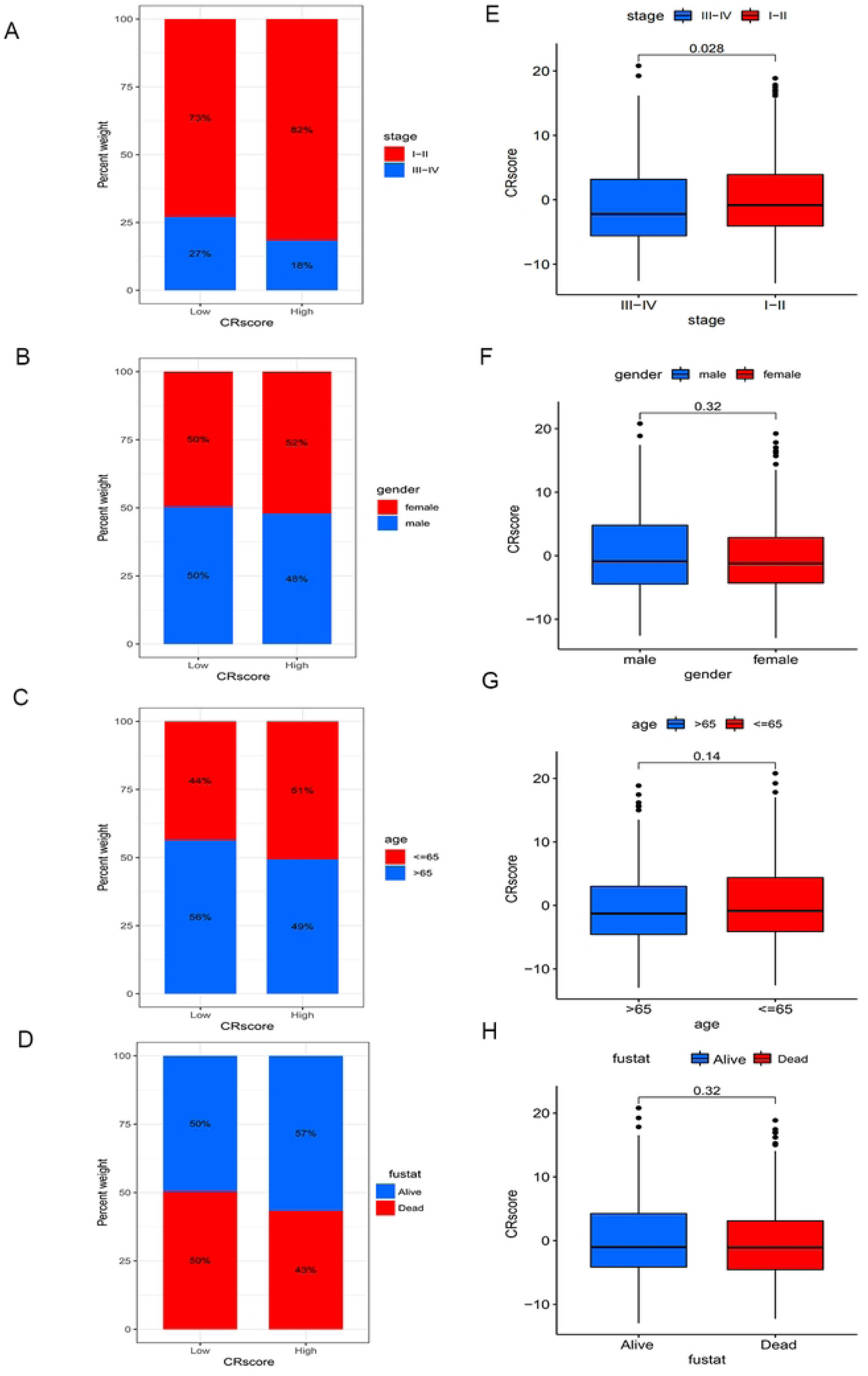
The relationship between CRscore and clinical features. **(A-D).** The abscissa represents CRscore type and the ordinate represents survival rate (The red areas represent phases **(A)**stage I-II **(B)** female **(C)** age<=65**(D)** dead and the blue areas represent phases **(A)**stage III-IV **(B)** male **(C)** age>65**(D)** alive**). (E-H).**The ordinate represents CRscore and the ordinate represents **(E)**stage **(F)** gender **(G)** age **(H)** fustat

**Fig 11.**
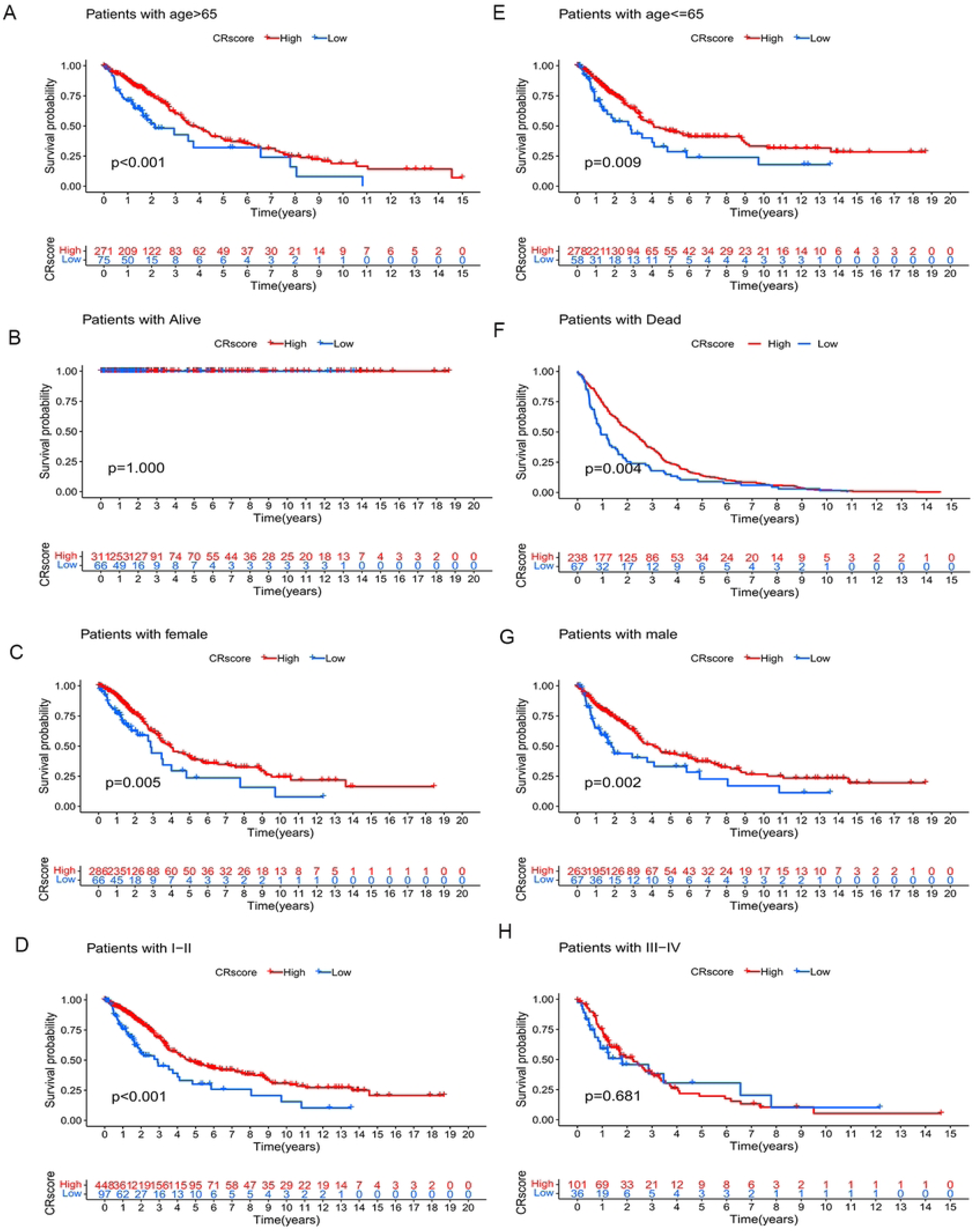
The relationship between CRcsore and survival of clinical features. **(A-H).** Kaplan-Meier curve was used to describe the difference in survival between groups with different clinical characteristics of high and low CRscore. The horizontal axis is survival time, and the vertical axis is **(A)**age<=65 **(B)**alive **(C)**female **(D)**stage I-II **(E)**age>65 **(F)**dead **(G)**male **(H)**stage III-IV.

### Immunotherapy for LUAD based on the CR score

Based on the CR score, we evaluated the differences in the expression of four common immune checkpoint proteins (PD1, PD-L1, PD-L2, and CTLA-4). The expression of immune checkpoint proteins was inversely related to high and low CR scores. In other words, the expression of immune checkpoint proteins was low in patients with high CR scores and high in patients with low CR scores (Figs. 12A–D). On the basis of the CR score, the IPS of LUAD was analyzed to predict its immunogenicity. Patients with high CR scores had higher IPS and IPS-CTLA4 scores (Figs. 12E–H). These results suggest that patients with LUAD with high CR scores may have a good response to CTLA4 immunotherapy (24). We analyzed the relationship between the CR score and the sensitivity of chemotherapy agents, which are used to treat LUAD, including cisplatin, gemcitabine, paclitaxel, vinorelbine and methotrexate. Patients with high CR scores were sensitive to cisplatin, gemcitabine, paclitaxel, and vinorelbine (P < 0.05) (Figs. 13A, C, D, E), and patients with low CR scores were sensitive to methotrexate (P < 0.05) (Fig. 13B). These results suggest that the CRscore tool is a dependable biological index of prognostic immunotherapy and clinical efficacy.

**Fig 12.**
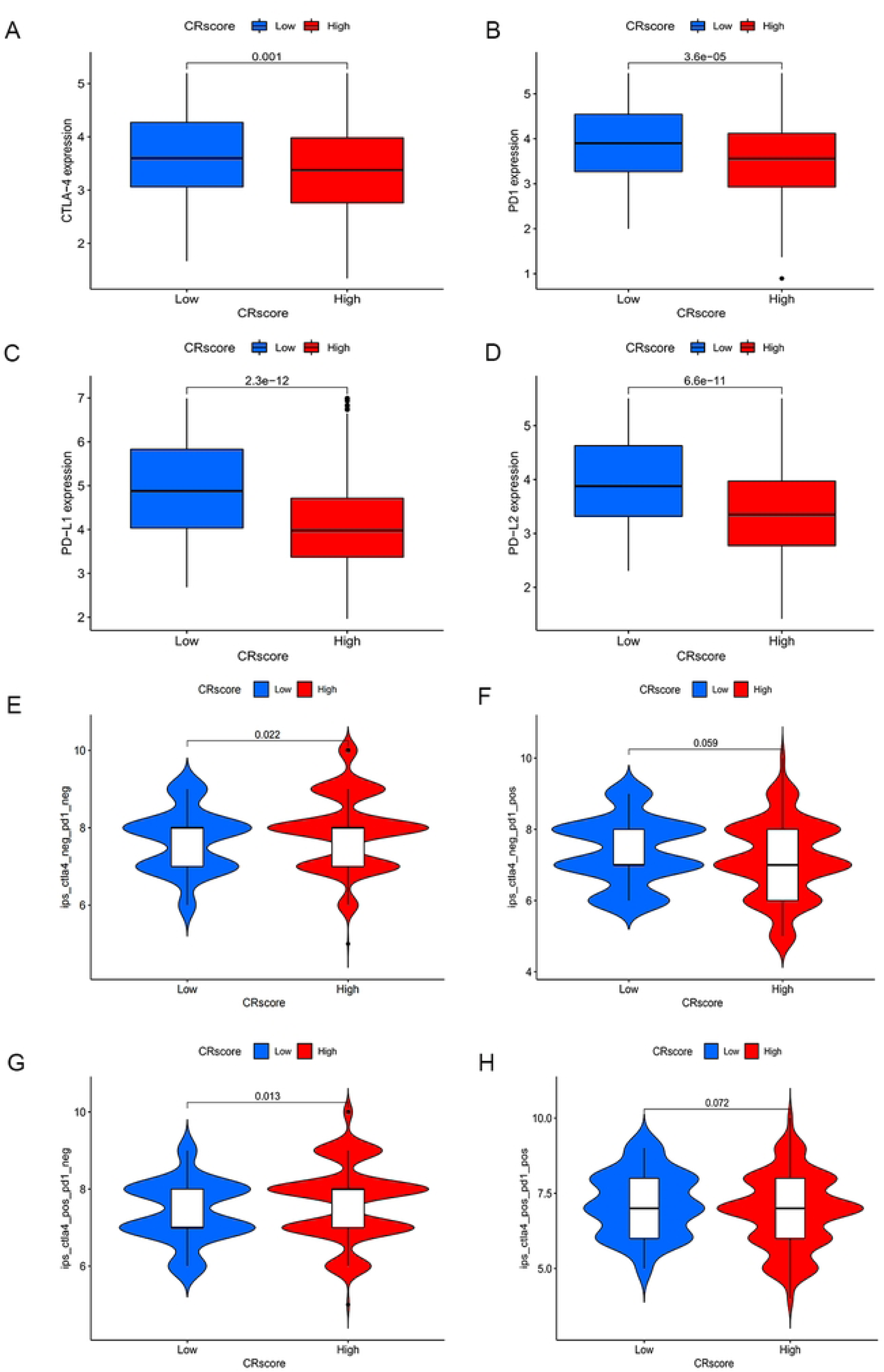
The relationship between CRcsore and immunotherapy. **(A-D).** Relationship between CRscore and immune checkpoints. The abscissa is CRscore, and the ordinate is the immune checkpoints **(A)** CTLA-4 **(B)** PD1 **(C)** PD-L1 **(D)** PD-L2 **(E-H).**Relationship between immunophenotypic score and high and low CRscore group. The abscissa is CRscore, and the ordinate is**(E)**ips_ctla4_neg_pd1_neg**(F)** ips_ctla4_neg_pd1_pos **(G)**ips_ctla4_pos_pd1_neg **(H)**ips_ctla4_pos_pd1_pos.

**Fig 13.**
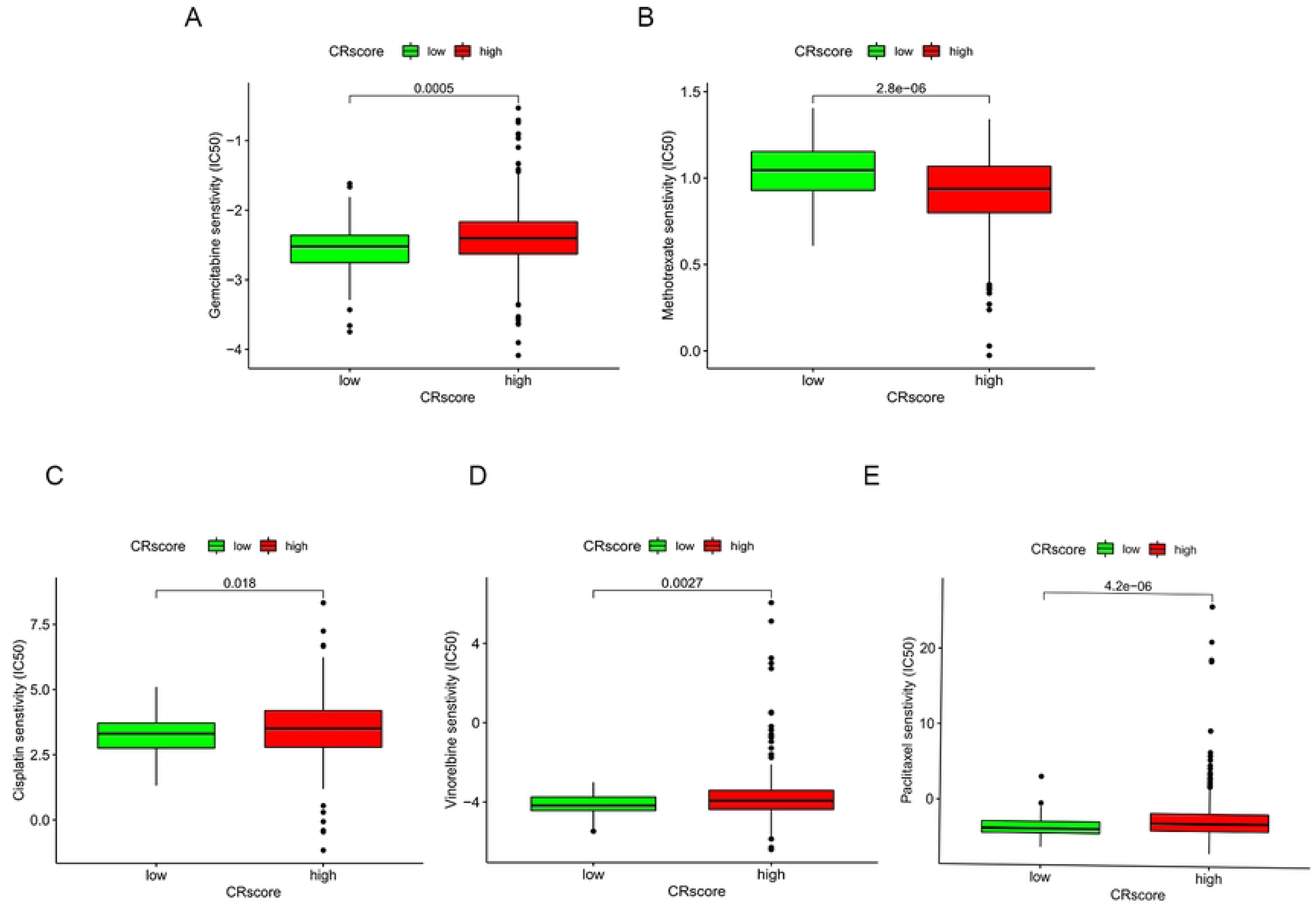
The relationship between CRscore and sensitivity to commonly used chemotherapeutic drugs. The abscissa is CRscore, and the ordinate is the sensitivity of chemotherapy drugs **(A)** gemcitabine **(B)** methotrexate **(C)** cisplatin **(D)** vinorelbine **(E)** paclitaxel.

### Clinical validation of CRRGs

To prove the accuracy of our CRscore tool and the strictness of the conclusions, we conducted a clinical trial on NPAS2, which is one of the CRRGs under study. We compared the expression of NPAS2 mRNA in LUAD tissues. NPAS2 was more highly expressed in LUAD tissues compared with healthy lung tissues according to qRT-PCR (P < 0.05) (Fig. 14C). Furthermore, we compared the expression of NPAS2 protein in LUAD tissues. As expected, Western blot showed that NPAS2 was significantly elevated in LUAD tissues compared with healthy lung tissues (P < 0.05) (Figs. 14A, B).

**Fig 14.**
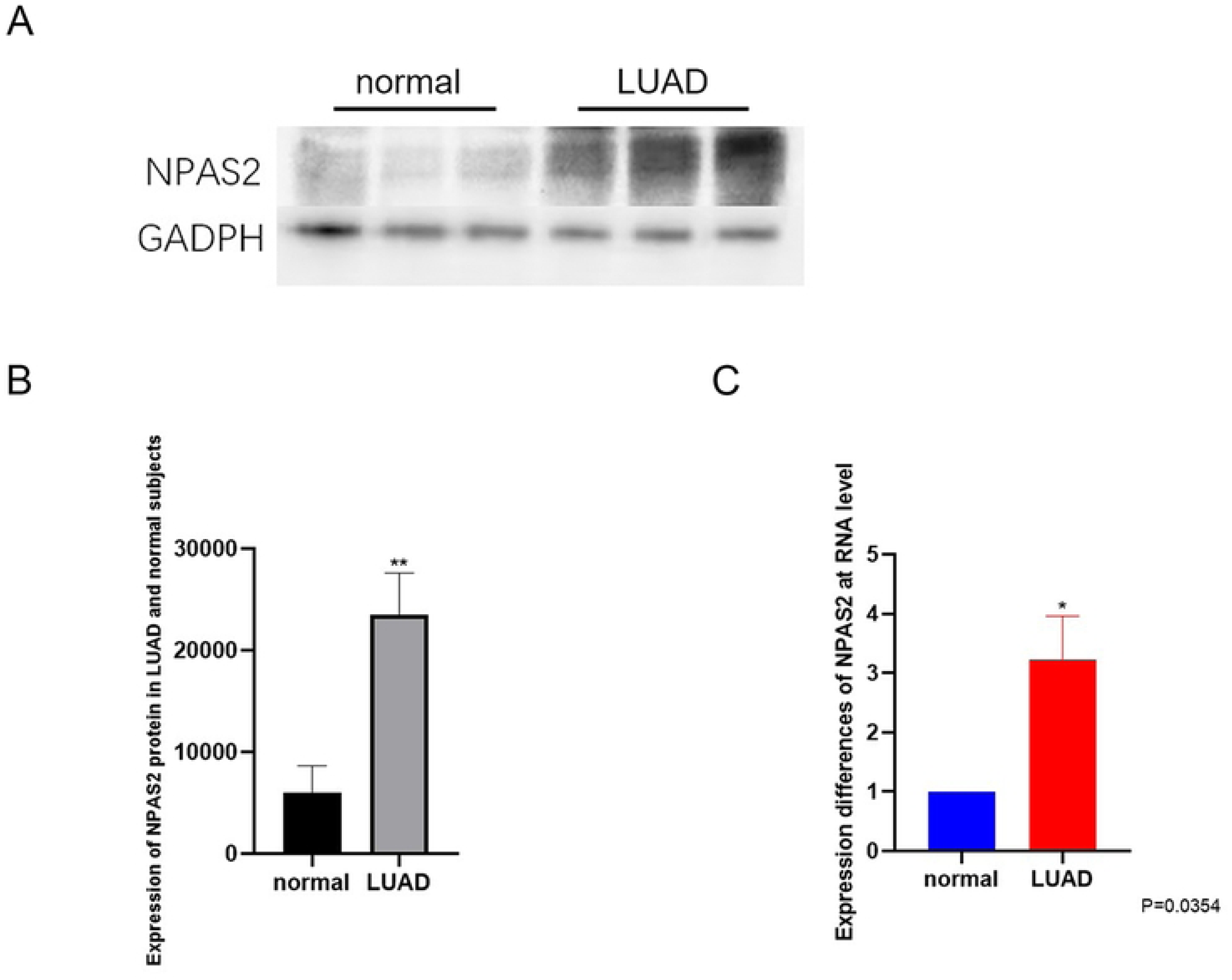
Clinical sample validation: **(A)** Western blot analysis of the influence of LUAD patients and normal patients on the expression level of NPAS2 protein. **(B)** Gray scanning quantitative analysis of protein. The mean of three independent groups was ± SD. The level of NPAS2 protein in LUAD patients was significantly different from that in normal patients (*P<0.05; ** P< 0.01) **(C)** differential expression of NPAS2 at RNA level between tumor patients and normal patients (* P<0.05).

## Discussion

The CR is a natural internal homeostatic mechanism that regulates the physiological light–dark cycle. Disruption of systemic and tissue-specific circadian mechanisms leads to changes in cell function, such as metabolism and cell division, both of which are highly associated with cancer (25). Pharmacological regulation of core CRRGs is a new approach for cancer treatment, and integrating circadian biology into cancer research offers new options for more effective cancer prevention, diagnosis, and treatment.

In this study, we found that the expression of CRRGs in samples from patients with LUAD was high under our preliminary exploration of the TCGA database. We speculate that CRRGs play an essential role in the occurrence and development of LUAD. We conducted an in-depth analysis of CRRDs in LUAD samples and established a CR scoring system (CRscore). We combined the CRscore tool with the expression of CRRGs, clinical features, TMB, and immune cell infiltration. As expected, the CR score was markedly associated with tumor mutations, immune cell infiltration, LUAD stage, and sex. Moreover, we showed that patients with higher CR scores had better survival. Encouragingly, the results were also tenable when we analyzed patients who have clinical characteristics uniformly to reduce the influence of other factors. The CR score was an independent prognostic factor according to the results of the univariate and multivariate Cox regression analyses.

Seven genes (NR1D2, AANAT, CRY1, NPAS2, CSNK1E, PER3, and CRY2) were significantly different between the LUAD group and the healthy group (P < 0.001). In the subsequent genotyping group of prognosis-associated CRRGs, the expression of these seven CRRGs was also significantly different among the three groups (P < 0.05). Therefore, we speculate that CRRGs play an essential role in the occurrence and development of LUAD. During the CR cluster analysis, the order of survival was CRcluster C > CRcluster B > CRcluster A. Coincidentally, the same conclusion was drawn among each CR gene cluster, as follows: gene cluster C > gene cluster B > gene cluster A . We hypothesized that the CR score was inextricably related to patient survival. This conjecture was confirmed in the subsequent analysis. As can be seen from the relationship between the CR score and the TMB, the CR score was positively correlated with the TMB. We speculated that high and low CR scores would reveal an anti-tumor process and tumor cell proliferation, respectively. A series of analyses on the gene population confirmed this speculation.

As can be seen from the survival analysis, the lower the CR score, the poorer the survival and the higher the tumor malignancy. Further, we detected four common immune checkpoint proteins and predicted their immunogenicity. We suggest that CTLA-4 immunotherapy is more suitable for patients with high CR scores. The sensitivity analysis of common chemotherapeutic drugs showed that most chemotherapeutic drugs were more effective in patients with high CR scores. From another perspective, we also know that the higher the CR score, the better the prognosis.

Although the CR score is a prognostic guide and a positive predictor of prognosis in patients with LUAD, some limitations of this study still need to be considered. First, the samples were obtained from public databases, which may have led to selection bias. Second, CRRGs in the database were transcribed from tumor tissues, making it improbable to recognize where the CRRGs identified in this study came from. Finally, not all patients with high CR scores will gain greater immunotherapy benefits, so more clinical factors need to be added to the prediction model to improve its accuracy.

In conclusion, we elucidated the significance of CRRGs in clinical practice, immune infiltration, and immunotherapy and gained several important insights. Our findings may guide the selection of combination strategies or lead to the manufacture of new immunotherapy drugs in the future. Our results provide new ideas for improving the clinical response of patients to immunotherapy, exploring new therapeutic targets, and promoting personalized cancer immunotherapy in the future.

## Ethical Statement

The authors are accountable for all aspects of the work in ensuring that questions related to the accuracy or integrity of any part of the work are appropriately investigated and resolved.

## Acknowledgments

Funding: This work was supported by the mechanism of curcumin ameliorating diabetic cardiomyopathy through RAGE mediated autophagy and its clinical application, led by Professor Kun Liu of affiliated hospital of Nantong University.(#MS12021070)

